# The ASCC2 CUE domain contacts adjacent ubiquitins to recognize K63-linked polyubiquitin

**DOI:** 10.1101/2021.10.17.464694

**Authors:** Patrick M. Lombardi, Sara Haile, Timur Rusanov, Rebecca Rodell, Rita Anoh, Julia G. Baer, Kate A. Burke, Lauren N. Gray, Abigail R. Hacker, Kayla R. Kebreau, Christine K. Ngandu, Hannah A. Orland, Emmanuella Osei-Asante, Dhane P. Schmelyun, Devin E. Shorb, Shaheer H. Syed, Julianna M. Veilleux, Ananya Majumdar, Nima Mosammaparast, Cynthia Wolberger

## Abstract

Alkylation of DNA and RNA is a potentially toxic lesion that can result in mutations and cell death. In response to alkylation damage, K63-linked polyubiquitin chains are assembled that localize the ALKBH3-ASCC repair complex to damage sites in the nucleus. The protein ASCC2, a subunit of the ASCC complex, selectively binds K63-linked polyubiquitin chains using its CUE domain, a type of ubiquitin-binding domain that typically binds monoubiquitin and does not discriminate among different polyubiquitin linkage types. We report here that the ASCC2 CUE domain selectively binds K63-linked diubiquitin by contacting both the distal and proximal ubiquitin. Whereas the ASCC2 CUE domain binds the distal ubiquitin in a manner similar to that reported for other CUE domains bound to a single ubiquitin, the contacts with the proximal ubiquitin are unique to ASCC2. The N-terminal portion of the ASCC2 α1 helix, including residues E467 and S470, contributes to the binding interaction with the proximal ubiquitin of K63-linked diubiquitin. Mutation of residues within the N-terminal portion of the ASCC2 α1 helix decreases ASCC2 recruitment in response to DNA alkylation, supporting the functional significance of these interactions during the alkylation damage response.

## Introduction

Ubiquitylation is a reversible, post-translational modification that regulates a vast array of cellular processes including proteasomal degradation, transcription, and the DNA damage response (1-3). The ubiquitin C-terminus is conjugated to protein substrates in a cascade of enzymatic reactions, most commonly forming a covalent linkage with the ε-amino group of a lysine side chain or the N-terminal α-amine (4). Ubiquitin itself can be ubiquitinated via one of its seven lysine residues or at its amino terminus, giving rise to homotypic or branched polyubiquitin chains with distinct topologies and biological functions (5). Different types of polyubiquitin chains are recognized by domains or motifs that bind specifically to the particular ubiquitin modification, thereby recruiting downstream effector proteins (6). In this manner, the diversity of ubiquitin signaling is predicated on the ability of ubiquitin-binding proteins to differentiate among the myriad types of polyubiquitin modifications present in the cell.

Lysine 63 (K63)-linked polyubiquitin chains play a non-degradative role in several DNA damage response pathways, including the response to DNA alkylation (2,7). The E3 ubiquitin ligase, RNF113A, assembles K63-linked polyubiquitin chains at sites of alkylation damage (7). These polyubiquitin chains recruit the ALKBH3-ASCC complex, which repairs the lesions (7). A subunit of the complex, ASCC2, binds to the K63-linked polyubiquitin chains via its CUE domain, a ubiquitin-binding domain of approximately 50 amino acids (Figure 1) (8,9). As shown for Cue1, Cue2, gp78, and Vps9, CUE domains bind the hydrophobic I44 patch of ubiquitin via conserved hydrophobic sequence motifs (Figure 1B) (8-11). These sequence motifs are conserved in ASCC2, suggesting that its CUE domain binds ubiquitin in a similar manner. Indeed, substitution of ASCC2 residue L506, which lies in the predicted ubiquitin-binding patch, abrogates ubiquitin binding *in vitro* and dramatically reduces formation of ASCC2 nuclear foci in response to alkylation damage (7). CUE domains typically make extensive interactions with a single ubiquitin within a polyubiquitin chain and exhibit little selectivity among different types of polyubiquitin chains (8-10,12). It is not known what other ASCC2 surfaces mediate interactions with ubiquitin and specify binding for K63-linked polyubiquitin.

**Figure 1.**
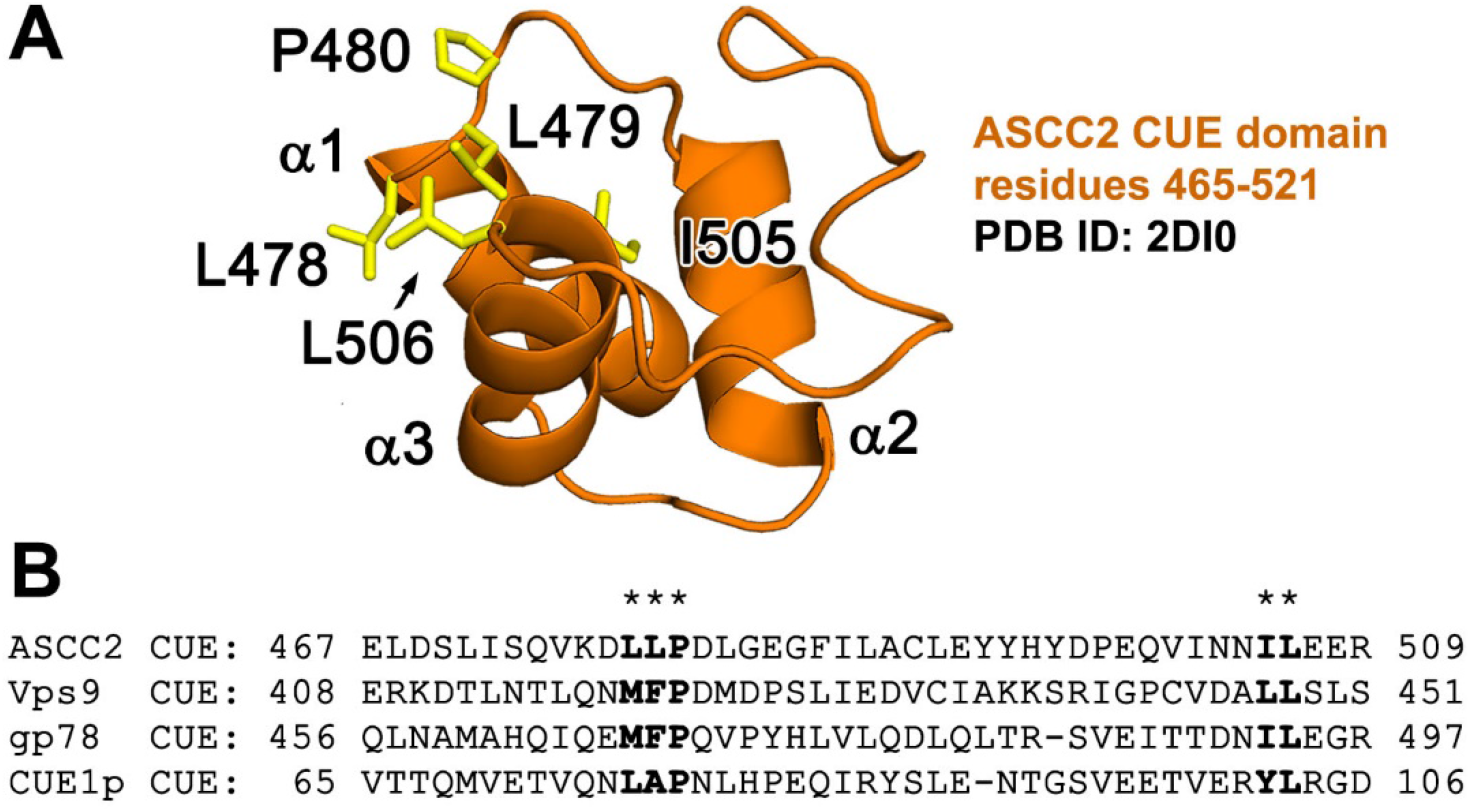
The ASCC2 CUE domain. (A) The ASCC2 CUE domain folds into a three-helix bundle. (B) CUE domains contain conserved sequence motifs on the α1 and α3 helices (in bold, below asterisks) that form a hydrophobic ubiquitin-binding surface (yellow sticks in figure 1A).

We report here that the ASCC2 CUE domain binds with higher affinity to K63-linked polyubiquitin as compared to monoubiquitin or other types of polyubiquitin chains. Using solution nuclear magnetic resonance (NMR), we show that a single ASCC2 CUE domain makes distinct contacts with the two adjacent ubiquitins within a single K63-linked polyubiquitin chain. In addition to mediating conserved interactions with the I44 patch of the distal ubiquitin, a separate region of the ASCC2 CUE domain forms additional interactions with the proximal ubiquitin in K63-linked polyubiquitin. Mutations in the ASCC2 CUE domain residues that contact the proximal ubiquitin disrupt recruitment of ASCC2 to repair foci, consistent with the importance of these residues in binding to K63-linked polyubiquitin. Together, our data show that the ASCC2 CUE domain makes multiple, linkage-specific interactions with adjacent ubiquitins to enhance the affinity of the ALKBH3-ASCC complex for K63-linked polyubiquitin chains at alkylation damage sites.

## Results

### The ASCC2 CUE domain has enhanced affinity and specificity for K63-linked polyubiquitin chains

The ASCC2 CUE domain comprises a three-helix bundle that spans residues 465-521 (Figure 1). The solution structure of the ASCC2 CUE domain has been deposited in the Protein Data Bank (PDB ID: 2DI0) (Figure 1B) and is similar to other experimentally determined CUE domain structures. The structure of the ASCC2 CUE domain superimposes with the Cue2 CUE domain (PDB ID: 1OTR) (8) with an RMSD of 0.92 Å over 37 alpha carbons and with the gp78 CUE domain (PDB ID: 2LVN) (10) with an RMSD of 1.34 Å over 41 alpha carbons. Like other CUE domains, ASCC2 has a cluster of hydrophobic residues on helix 1 and on helix 3, which are predicted to bind to the I44 patch of ubiquitin as in previously characterized CUE:ubiquitin interactions (8-12) (Figure 1B). The previous finding that a substitution at L506 in helix 3 leads to defects in ASCC2 recruitment in cells (7) is consistent with a role for this surface in ASCC2 binding to ubiquitin.

In order to determine whether the ASCC2 CUE domain binds in a similar manner to ubiquitin irrespective of whether it is incorporated into a polyubiquitin chain, we used isothermal titration calorimetry (ITC) to compare binding of ASCC2 CUE domain constructs to monoubiquitin and to K63-linked diubiquitin (K63Ub_2_). As shown in Figures 2A and 2B, we found that the ASCC2 CUE domain binds with lower affinity to monoubiquitin than to K63Ub_2_. The equilibrium dissociation constant (*K*_d_) for monoubiquitin was 57.1 μM ± 5.0 μM (Figure 2A), while ASCC2 bound much more tightly to K63Ub_2_, with a *K*_d_ of 8.7 μM – 10.4 μM (Figure 2B). The affinity of the isolated ASCC2 CUE domain for K63Ub_2_ is similar to that of full-length ASCC2, which binds to K63Ub_2_ with a *K*_d_ of 8.8 μM ± 0.9 μM (Figure 2C). The similar equilibrium dissociation constants suggest that the majority of the affinity comes from the interaction between the CUE domain and K63Ub_2_. Importantly, the approximately 4 – 7-fold enhancement of ASCC2 CUE domain affinity for K63Ub_2_ as compared to monoubiquitin is higher than the 1.0 – 1.8-fold enhancement in affinity that has been reported for other CUE domains binding to polyubiquitin versus monoubiquitin (10,11).

**Figure 2.**
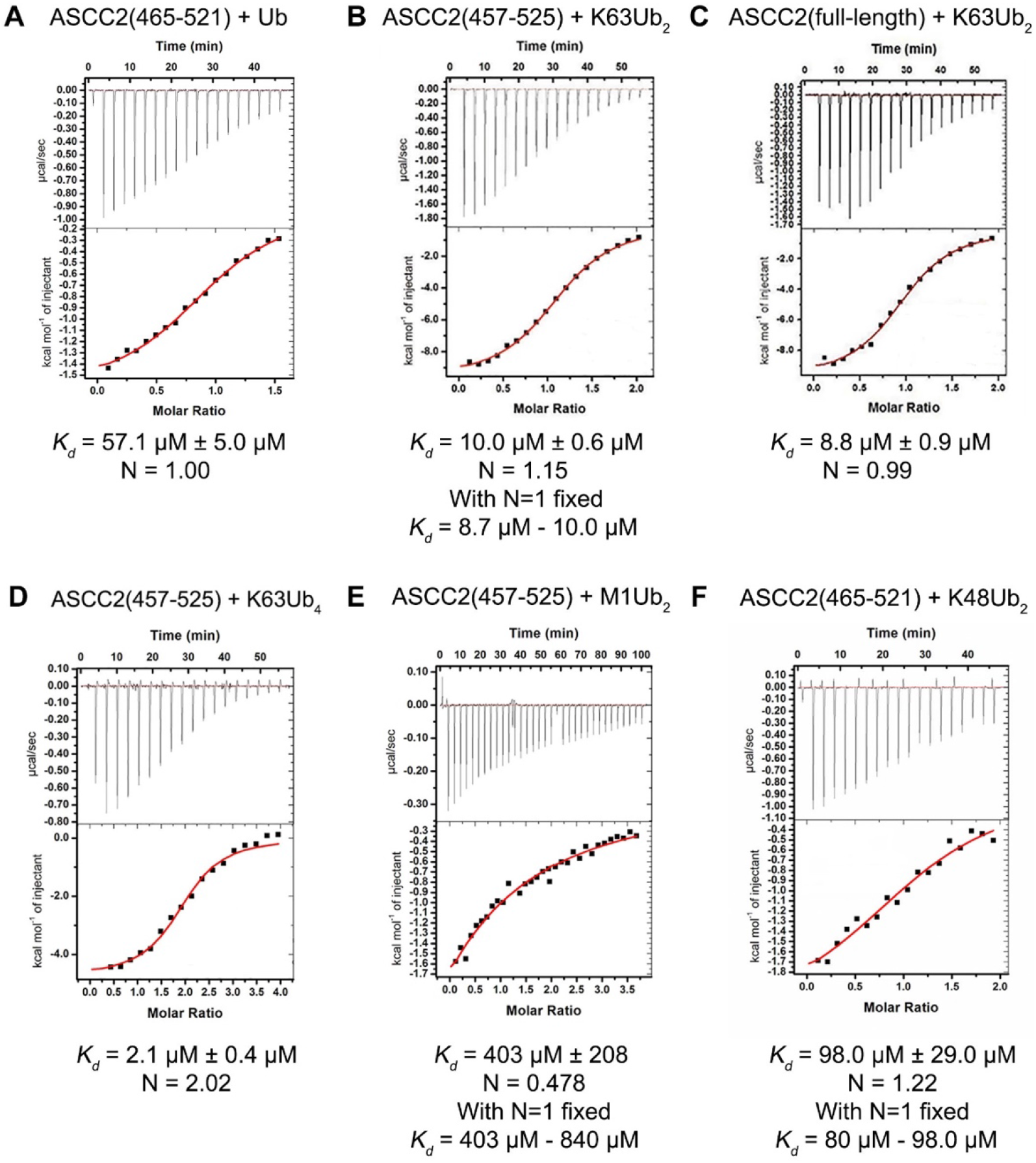
The ASCC2 CUE domain binds K63Ub_2_ with enhanced affinity and 1:1 stoichiometry. (A-C) ITC data show that full-length ASCC2 and isolated ASCC2 CUE domain bind K63Ub_2_ with greater affinity than monoubiquitin (Ub). (D) Two ASCC2 CUE domains bind per K63-linked tetraubiquitin chain (K63Ub_4_). This 1:1 ASCC2 CUE domain to K63Ub_2_ ratio is observed in Figures 2B and 2C as well. (E) The ASCC2 CUE domain does not exhibit enhanced binding affinity for linear diubiquitin (M1Ub_2_) or (F) K48-linked diubiquitin (K48Ub_2_).

Since very weak binding is difficult to measure accurately by ITC, we also estimated the affinity of the ASCC2 CUE domain for monoubiquitin using nuclear magnetic resonance (NMR) spectroscopy. The ^1^H,^15^N-HSQC spectra of ^15^N-labeled ASCC2(465-521) were recorded in the presence of increasing amounts of monoubiquitin and the chemical shift perturbations (CSPs) for four ASCC2 residues at the ubiquitin-binding interface were used to calculate the average *K*_d_ value (Supplementary Figure 1) (13). The *K*_d_ value of 39.6 μM ± 1.6 μM determined by NMR suggests somewhat tighter monoubiquitin binding than the *K*_d_ determined by ITC (57.1 μM ± 5.0 μM Figure 2A), although still substantially weaker than that measured for K63Ub_2_ (*K*_d_ = 8.7 μM – 10.4 μM Figure 2B).

The stoichiometry of the ASCC2 CUE domain binding K63-linked polyubiquitin chains is also different from that of previously studied CUE domains. CUE domains from other ubiquitin-binding proteins, such as gp78, bind diubiquitin with a ratio of two CUE domains per diubiquitin (10), indicating that each CUE domain binds to one ubiquitin in the diubiquitin chain. The ASCC2 CUE domain, however, binds K63Ub_2_ in a 1:1 ratio (Figure 2B), and this ratio is conserved in the binding of full-length ASCC2 to K63Ub_2_ (Figures 2C). Importantly, the observed molar ratio of one ASCC2 CUE domain per K63Ub_2_ is preserved in the context of longer polyubiquitin chains. As shown in Figure 2D, the ASCC2 CUE domain binds to K63-linked tetraubiquitin with a molar ratio of 2:1, consistent with each CUE domain binding to two ubiquitins within the tetraubiquitin chain. Interestingly, the affinity of the ASCC2 CUE domain for K63-linked tetraubiquitin is about 4-fold higher than its affinity for diubiquitin (Figures 2B and 2D).

In order to test the specificity of ASCC2 for K63-linked diubiquitin as compared to other linkage types, we measured the *K*_d_ of the ASCC2 CUE domain for linear and K48-linked diubiquitin (K48Ub_2_). The affinity of the ASCC2 CUE domain for linear diubiquitin (M1Ub_2_) was extremely weak, with a *K*_*d*_ of about 400 μM (Figure 2E). This result was surprising given that linear and K63-linked polyubiquitin adopt a similar extended topology (14,15). The affinity of the ASCC2 CUE domain for K48Ub_2_ (Figure 2F), with a *K*_*d*_ of about 98 μM, was similar to that measured for monoubiquitin. These results indicate that the enhanced binding affinity of the ASCC2 CUE domain for polyubiquitin compared to monoubiquitin is specific to K63-linked chains.

### The ASCC2 CUE domain forms different contacts with the distal and proximal ubiquitin of K63Ub_2_

The higher affinity of ASCC2 for di- or tetra-ubiquitin and the molar ratio of one ACSS2 CUE domain per diubiquitin are consistent with a single CUE domain simultaneously contacting the linked proximal and distal ubiquitin. Given the small size and asymmetric fold of the CUE domain, ASCC2 would need to form different binding interfaces with the two ubiquitin monomers. We used NMR chemical shift mapping experiments to compare ASCC2 CUE domain contacts with the distal and proximal ubiquitins of K63Ub_2_. In order to distinguish the two covalently linked ubiquitin monomers, we generated diubiquitin with either the proximal or the distal ubiquitin isotopically labelled with ^15^N. The ^1^H,^15^N-HSQC spectra of the differently labeled K63Ub_2_ were recorded in the presence of increasing concentrations of the ASCC2 CUE domain (Figure 3). The CSP values for the ^15^N-labeled distal ubiquitin of K63Ub_2_ titrated with the ASCC2 CUE domain (Figure 3A) are similar to those reported for other CUE domains interacting with monoubiquitin (10,11). Common features include relatively large CSP values for residues in and around the I44 patch, such as R42, I44, G47, and K48, and for residues 70-74 at the C-terminal tail of ubiquitin (Figure 3A). The similar CSP values suggest the distal ubiquitin of K63Ub_2_ binds the ASCC2 CUE domain using the same surface as previously reported CUE:ubiquitin interactions (10,11).

**Figure 3.**
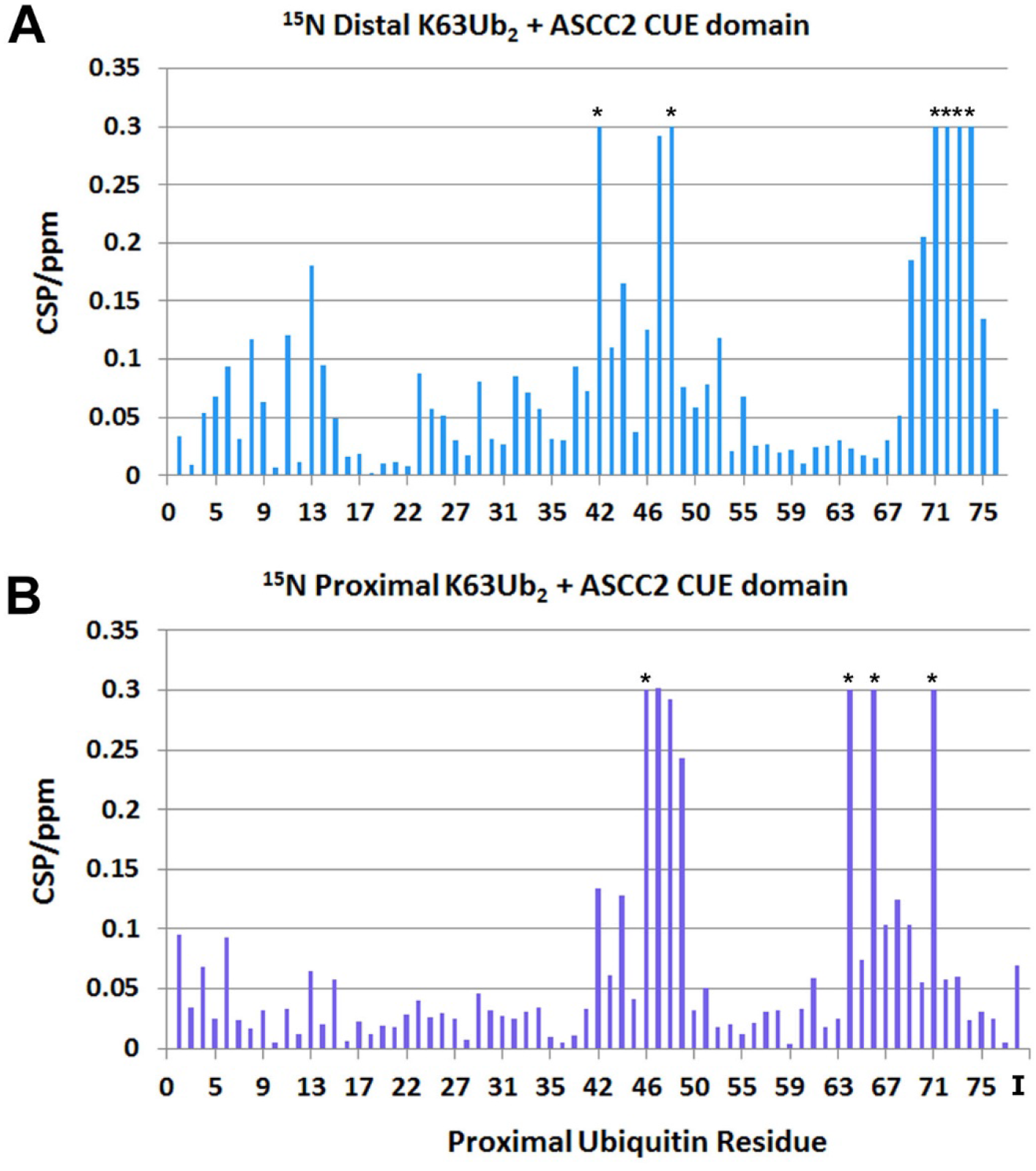
The ASCC2 CUE domain makes different interactions with the distal and proximal ubiquitins of K63Ub_2_. Differences in the CSP values recorded for the ^15^N-labeled distal (A) and proximal (B) ubiquitins of K63Ub_2_ titrated with the ASCC2 CUE domain suggest that ASCC2 makes different contacts with the distal and proximal ubiquitin. An asterisk (*) indicates the resonance disappeared during the titration. The “I” in Figure 3B denotes the K63Ub_2_ isopeptide bond.

The CSP values for the ^15^N-labeled proximal ubiquitin of K63Ub_2_ titrated with ASCC2(465-521) (Figure 3B), however, were markedly different from those observed for the ^15^N-labeled distal ubiquitin. The CSP values for ubiquitin residues V70, R72, L73, and R74 were smaller in the experiments with ^15^N-labeled proximal ubiquitin (Figure 3B) as compared to the experiments with ^15^N-labeled distal ubiquitin (Figure 3A). In addition, large CSP values for residues E64 and T66 (Figures 3B) were unique to the proximal ubiquitin. The large CSP values for proximal ubiquitin residues E64 and T66 and the small CSP values for residues in the ubiquitin C-terminal tail suggest that the ASCC2 CUE domain contacts the proximal ubiquitin in a non-canonical manner. The differences between the proximal and distal ubiquitin spectra suggest that the two ubiquitins form different interactions with the ASCC2 CUE domain.

To determine the contribution of proximal ubiquitin residues E64 and T66 to ASCC2 CUE domain binding, we measured the affinity of the ASCC2 CUE domain for K63Ub_2_ bearing side chains substitutions at proximal ubiquitin residues E64 and T66 (K63Ub_2_ E64A/T66A_prox_) using ITC. As shown in Figure 4, the ASCC2 CUE domain binds K63Ub_2_ E64A/T66A_prox_ with a *K*_d_ in the range of 45.9 μM – 90.9 μM, approximately 3.5 – 7.0 times more weakly than wild-type K63Ub_2_. This result supports the NMR data (Figure 3B) in suggesting that the ASCC2 CUE domain interacts with residues E64 and T66 of the proximal ubiquitin in K63Ub_2_.

**Figure 4.**
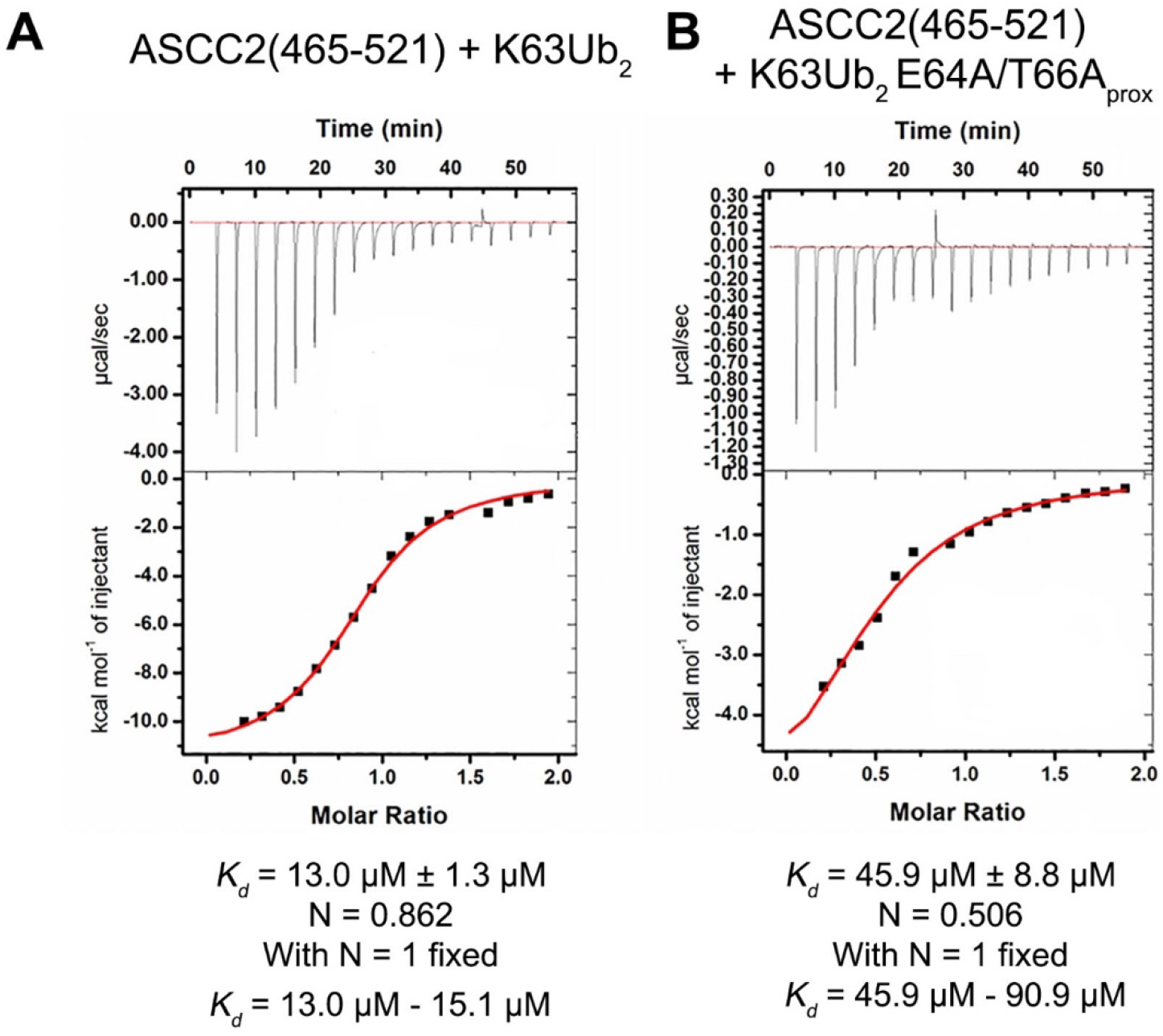
The ASCC2 CUE domain binds K63Ub_2_ E64A/T66A_prox_ with reduced affinity relative to wild-type K63Ub_2_. ITC data show that ASCC2(465-521) binds wild-type K63Ub_2_ (A) with ∼3.5-7.0X greater affinity than K63Ub_2_ E64A/T66A_prox_ (B).

### The N-terminal portion of the ASCC2 α1 helix is important for K63-linked polyubiquitin binding and recruitment to DNA damage foci

To determine which ASCC2 residues interact with K63-linked polyubiquitin, the ^1^H,^15^N-HSQC spectra of ^15^N-labeled ASCC2 CUE domain were recorded in the presence of increasing concentrations of monoubiquitin and of K63Ub_2_ (Figures 5). ASCC2 residues L479 and L506, from the conserved CUE domain hydrophobic sequence motifs in the α1 and α3 helices, respectively, exhibited large CSP values when the ASCC2 CUE domain was titrated with monoubiquitin or K63Ub_2_ (Figures 5). This finding suggests that, as with other CUE domains (8-12), the hydrophobic surface created by the conserved sequence motifs is the binding site for monoubiquitin and one of the ubiquitins in K63Ub_2_. The distal ubiquitin of K63Ub_2_ is most likely to bind the hydrophobic surface formed by the conserved sequence motifs given the similarities between its CSP values (Figure 3A) and those reported for monoubiquitin titrated with other CUE domains (10,11).

**Figure 5.**
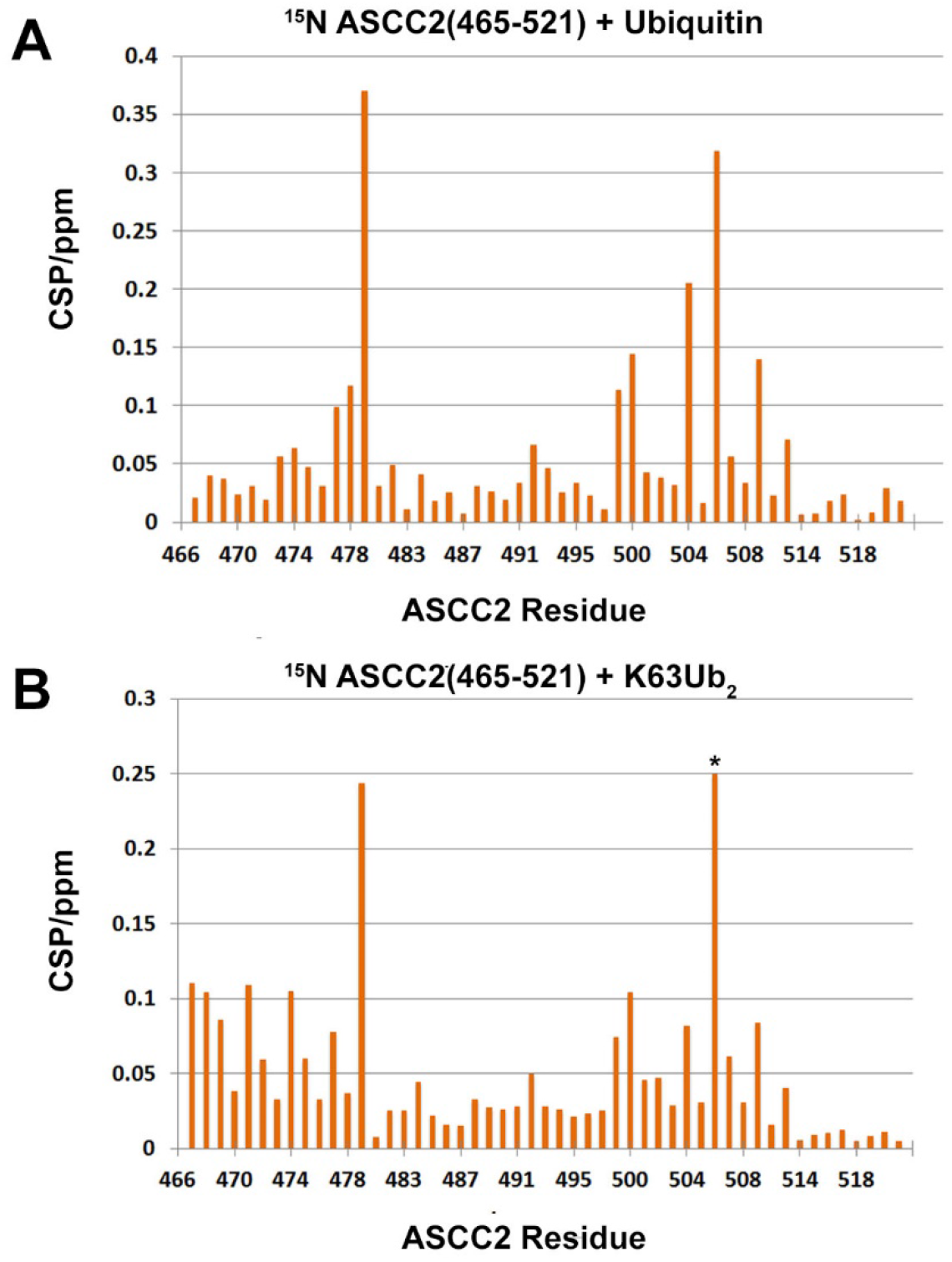
The ASCC2 CUE domain utilizes conserved sequences from the α1 and α3 helices, along with the N-terminal end of the α1 helix, to bind K63-linked polyubiquitin chains. CSP values recorded from the ^1^H-^15^N-HSQC spectra of ^15^N-labeled ASCC2 CUE domain titrated with monoubiquitin (A) and K63Ub_2_ (B) differ most substantially from residues 466-474, suggesting that the N-terminal portion of the ASCC2 α1 helix may participate in binding to K63-linked polyubiquitin chains. An asterisk (*) indicates the resonance disappeared during the titration.

The ASCC2 residues that interact with the proximal ubiquitin of K63-linked diubiquitin would be expected to have larger CSP values when the CUE domain is titrated with K63Ub_2_ than when it is titrated with monoubiquitin. The α1 helix (residues 465-479; Figure 1A) is the only region of the ASCC2 CUE domain that has dramatically different CSP values in the presence of K63Ub_2_ compared to monoubiquitin (Figure 5). The majority of residues from the N-terminal end of the α1 helix have relatively small CSP values when titrated with monoubiquitin (Figure 5A). When titrated with K63Ub_2_, however, the CSP values for residues from the N-terminal end of the α1 helix are larger (Figure 5B). The increased CSP values for residues at the N-terminal end of the α1 helix in the presence of K63Ub_2_ compared to monoubiquitin suggest that these residues may bind the proximal ubiquitin of K63Ub_2_.

To test the hypothesis that ASCC2 residues at the N-terminal end of the α1 helix bind the proximal ubiquitin of K63Ub_2_, we made point mutations at ASCC2 residues E467, S470, and L471 and assayed their effects on ASCC2 CUE domain binding to K63Ub_2_. While ITC experiments showed that the affinity of the ASCC2(465-521) L471A mutant for K63Ub_2_ was nearly identical to that of wild-type ASCC2(465-521) (Figure 6A), the ASCC2 E467A mutant bound 3.6 – 5.0-fold more weakly, with a *K*_d_ in the range of 46.9 μM - 65.4 μM (Figure 6B). The ASCC2 S470R mutant bound with even lower affinity, with an apparent *K*_d_ of 90.9 μM ± 23.1 μM (Figure 6C). An ASCC2 E467R/S470R double mutant bound K63Ub_2_ with an apparent *K*_d_ of 92.6 μM ± 20.9 μM (Figure 6D). The decrease in K63Ub_2_ binding affinity observed upon mutating the α1 helix stands in contrast to the effect observed upon altering other ASCC2 CUE domain regions that could potentially interact with the proximal ubiquitin of K63Ub_2_, such as the α2 helix, or the loop connecting the α2 and α3 helices, where mutations resulted in little change in binding affinity (Supplementary Figure 2). The decreased affinity observed for the E467 and S470 mutant proteins is consistent with ^1^H, ^15^N-HSQC data (Figure 5) suggesting that the ASCC2 CUE domain binds the proximal ubiquitin of K63Ub_2_ using a second, previously uncharacterized, interaction site located at the N-terminal end of the α1 helix. The presence of a second binding site on the ASCC2 CUE domain could account for the enhanced affinity of the ASCC2 CUE domain for K63Ub_2_ relative to monoubiquitin and the 1:1 stoichiometry of ASCC2 CUE:K63Ub_2_ binding (Figure 2).

**Figure 6.**
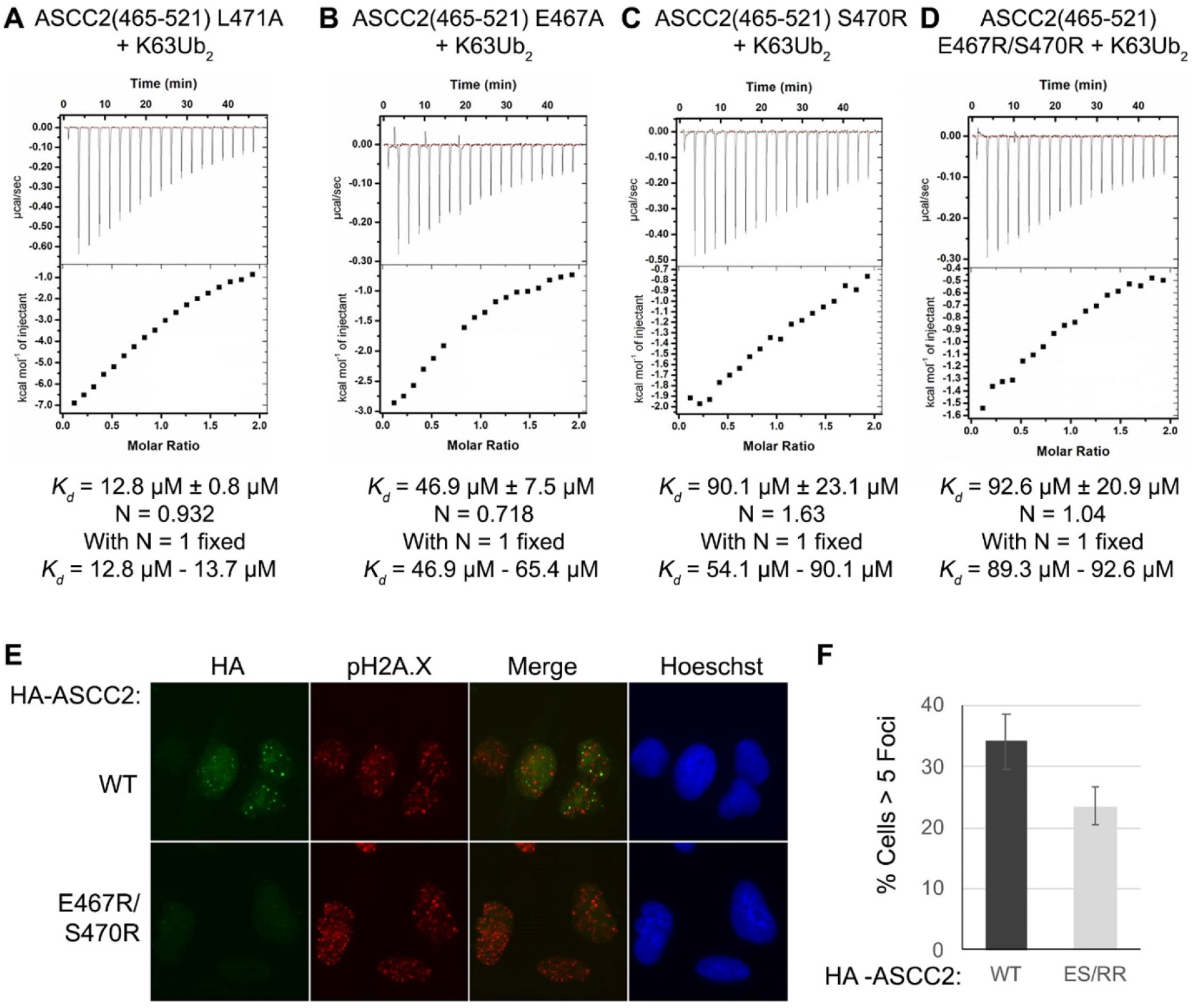
ASCC2 residues E467 and S470 bind the proximal ubiquitin of K63Ub_2_. (A) The ASCC2(465-521) L471A mutant binds K63Ub_2_ with nearly the same affinity as wild-type ASCC2(465-521) (Figure 4A). (B) The ASCC2(465-521) E467A mutant, however, binds K63Ub_2_ approximately 3.6 – 5.0-fold more weakly than wild-type ASCC2(465-521) and (C) the ASCC2(465-521) S470R mutant and (D) the ASCC2(465-521) E467R/S470R double mutant bind approximately 7.0-fold more weakly. These results suggest that ASCC2 residues E467 and S470 participate in the binding interaction with K63-linked polyubiquitin chains. (E) HA-tagged ASCC2, or the E467R/S470R mutant, were expressed in U2OS cells, then treated with 0.5 mM MMS for six hours. Immunofluorescence for HA and pH2A.X were performed after extraction with Triton X-100, as shown, with Hoechst used as the nuclear counter stain. (F) Quantification of foci formation. N = 3 replicates and error bars indicate +/- S.D. of the mean.

To test the functional importance of the N-terminal portion of the ASCC2 CUE domain α1 helix in cells, we studied the effect of an E467R/S470R double mutation on recruitment of ASCC2 to alkylation damage-induced foci. As compared to wild-type ASCC2, the E467R/S470R double mutant had significantly reduced ASCC2 foci after alkylation damage was induced with methyl methanesulphonate (MMS) (Figures 6E and 6F). This mutant was expressed at levels similar to the wild-type protein, suggesting that the defect is not due to a loss of expression due to misfolding or other global defect (Supplementary Figure 3). These results are consistent with a role for the N-terminal portion of the ASCC2 CUE domain α1 helix in its recruitment during the DNA damage response.

### Model of the ASCC2 CUE domain binding to K63-linked diubiquitin

We modeled the interaction between the proximal ubiquitin of K63Ub_2_ and the ASCC2 CUE domain using the HADDOCK protein-docking server (16,17). We first generated a model of the interaction between the ASCC2 CUE domain and the distal ubiquitin based on the gp78 CUE:monoubiquitin complex (PDB ID: 2LVO) (10) by superimposing residues 465-521 of the ASCC2 CUE domain structure (PDB ID: 2DI0) on the gp78 CUE domain. Given the similarity between the CSP values for the distal ubiquitin of K63Ub_2_ titrated with the ASCC2 CUE domain (Figure 3A) and the CSP values for monoubiquitin titrated with the gp78 CUE domain (10), it is likely that these interactions are structurally similar. Distance restraints based on NMR CSP data and mutagenesis data were utilized by the HADDOCK server to guide the docking of the proximal ubiquitin of K63Ub_2_ to the ASCC2 CUE domain. ASCC2 residues E467 and S470, and proximal ubiquitin residues E64 and T66, were specified as residues likely to be at the binding interface based on the deleterious effect of substitutions at these residues on binding (Figures 4B and 6B-D). In addition, proximal ubiquitin residues with CSP values greater than 2σ were also specified as likely to be at the interface with ASCC2. These proximal ubiquitin residues include A46, G47, K48, Q49, and L71. The resulting model of the ASCC2(465-521):K63Ub_2_ complex is shown in Figure 7.

**Figure 7.**
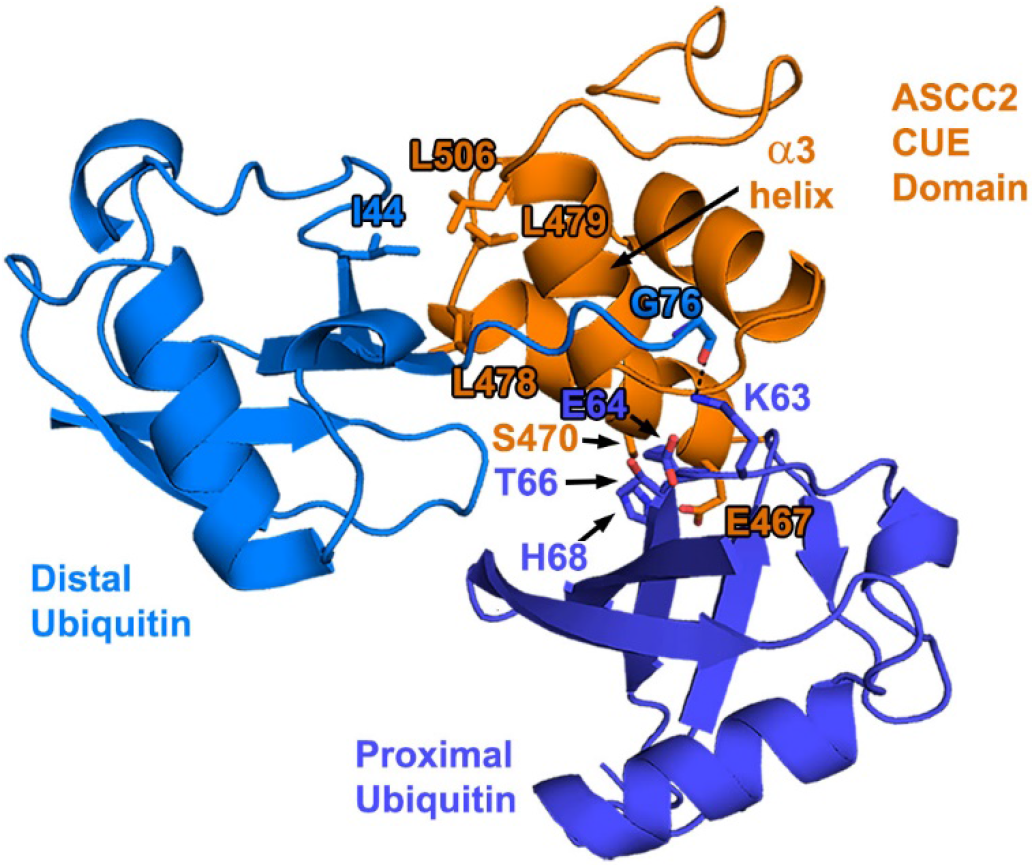
Model of the ASCC2(465-521):K63Ub_2_ complex. The ASCC2 CUE domain was superimposed onto the gp78 CUE domain structure in the gp78 CUE:ubiquitin complex (PDB ID: 2LVO). The proximal ubiquitin of K63Ub_2_ was docked to the complex using the HADDOCK server. In this model, binding of the I44 patch of the distal ubiquitin to the hydrophobic α1 and α3 sequence motifs of ASCC2, mediated by residues including L479 and L506, positions the C-terminal tail of the distal ubiquitin to interact with the α3 helix of ASCC2. Isopeptide bond formation at proximal ubiquitin residue K63 (dashed line), places adjacent proximal ubiquitin residues E64 and T66 within binding distance of the ASCC2 CUE domain. The proximal ubiquitin of K63Ub_2_ docks against the N-terminal end of the CUE domain α1 helix, where ASCC2 residues E467 and S470 interact with proximal ubiquitin residues.

## Discussion

The preferential binding of ASCC2 to K63-linked polyubiquitin chains stands in contrast to other CUE domain proteins, which show more modest enhancement of binding to mono-versus polyubiquitin and little selectivity among polyubiquitin chain types (10,12). We found that the affinity of the ASCC2 CUE domain for K63Ub_2_ is approximately 4 – 7-fold stronger than its affinity for monoubiquitin and that the ASCC2 CUE domain binds to di- and tetraubiquitin in a ratio of one CUE domain per diubiquitin (Figure 2). While interactions with the distal ubiquitin are similar to those observed for other CUE domains bound to monoubiquitin (8-12), ASCC2 contacts the proximal ubiquitin in a non-canonical manner using residues from the N-terminal portion of the α1 helix (Figures 5 & 6). By contrast, CUE domains from other proteins, such as Cue1 and gp78, make fewer contacts with adjacent ubiquitins within polyubiquitin chains and, accordingly, exhibit smaller enhancements in their affinities for polyubiquitin chains compared to monoubiquitin (10,11). For example, the CUE domain from the protein Cue1 binds to the I44 patch of the proximal ubiquitin in K48Ub_2_ while also contacting G75 and the C-terminus of the distal ubiquitin (11). The *K*_d_ for the Cue1 CUE domain binding K48Ub_2_ is 95 μM, compared to 173 μM for binding to monoubiquitin (11). This enhancement is much more modest than the 4 – 7-fold enhancement observed for the ASCC2 CUE domain. The gp78 CUE domain contacts T66 of the proximal ubiquitin in K48Ub_2_ while binding the I44 patch of the distal ubiquitin, resulting in an enhancement in affinity for the distal ubiquitin in K48Ub_2_ relative to the proximal ubiquitin (10). Overall, however, the gp78 CUE domain binds K48Ub_2_ and monoubiquitin with virtually equal affinity of about 12.4 μM and 12.8 μM, respectively (10). Despite the modest enhancement in affinity for polyubiquitin chains exhibited by the gp78 and Cue1 CUE domains, interacting with adjacent ubiquitins simultaneously is important for their biological functions. For these CUE domains, the interactions described above properly position ubiquitin ligases to add to growing polyubiquitin chains (10,11). For the ASCC2 CUE domain, interactions with adjacent ubiquitins strengthen the affinity for ASCC2’s biologically relevant target, K63-linked polyubiquitin chains (Figure 2), and increase the recruitment of the ALKBH3-ASCC repair complex to alkylation damage sites (Figures 6E-F).

While the ASCC2 CUE domain has been shown to bind K63Ub_2_ with enhanced affinity relative to M1Ub_2_ and K48Ub_2_, the structural basis for this selectivity has not been fully elucidated. Linear polyubiquitin chains, and possibly K48-linked polyubiquitin chains, can adopt similar conformations to the K63-linked polyubiquitin chain shown in Figure 7 (14,15), and interact with the same surfaces of the ASCC2 CUE domain. The affinity of the ASCC2 CUE domain for M1Ub_2_ and K48Ub_2_ relative to K63Ub_2_ is much weaker, however (Figure 2). We speculate that ASCC2 CUE domain interactions with proximal ubiquitin residues near the K63 isopeptide linkage, including residues E64 and T66, contribute to the selectivity for K63-linked polyubiquitin. Consistent with this, we found that alanine substitutions of proximal ubiquitin residues E64 and T66 reduced the affinity for K63Ub_2_ by about 3.5 – 7.0-fold (Figure 4). Furthermore, it is not known which proximal ubiquitin surface in K63Ub_2_ interacts with the N-terminal portion of the ASCC2 CUE domain α1 helix and whether the proximal ubiquitins of M1Ub_2_ and K48Ub_2_ are capable of making similar contacts. Elucidating the structural details of the interactions between K63Ub_2_ residues E64, T66, and the proximal ubiquitin as a whole, with the ASCC2 CUE domain will be the subject of continued investigation of the basis for ASCC2 domain selectivity.

## Experimental Procedures

### Plasmids for protein expression

ASCC2 constructs containing amino acids 457-525 or 465-521 were inserted into a pPROEX HTa vector with an N-terminal polyhistidine tag followed by a tobacco etch virus (TEV) protease recognition sequence. Full-length ASCC2 was inserted into the pET28 vector. Wild-type ubiquitin residues 1-76 (wt Ub), along with mutant ubiquitin constructs containing K48R/K63R substitutions (K48R/K63R Ub) or a D77 extension (D77 Ub), were inserted into the pET-3a vector.

### Plasmids for cell-based studies

Full-length human ASCC2 cDNA cloned into pENTR-3C and pET-28a-Flag was previously described (7). The ASCC2 E467R/S470R mutant cDNA was synthesized (IDT), cloned into pENTR-3C and pET-28a-Flag, and confirmed by Sanger sequencing. For human cell expression, ASCC2 E467R/S470R was subcloned into pHAGE-CMV-3XHA using Gateway recombination.

### Expression and purification of the ASCC2 CUE domain

BL21 DE3 *E. coli* cells were transformed with pPROEX HTa vector containing the ASCC2 constructs, plated on LB agar containing 100 μg/mL ampicillin, and incubated overnight at 37°C. Single colonies were used to inoculate 5-mL aliquots of LB media with 100 μg/mL ampicillin. The 5-mL cultures were grown overnight at 37°C with 250 rpm shaking until saturation. The 5-mL colonies were used to inoculate 1-L cultures of LB media with 100 μg/mL ampicillin, which were grown at 37°C with 250 rpm shaking until reaching an OD_600_ between 0.5 and 0.8. Protein expression was induced by adding 0.5 mM IPTG, and allowed to continue overnight at 16°C with 250 rpm shaking. Following protein expression, cell were pelleted by centrifugation, resuspended in a buffer consisting of 50 mM Tris pH 7.5 and 1 mM PMSF, and lysed by sonication on ice. Cell lysate was centrifuged at 17,500 x *g* for 30 minutes at 4°C, passed through a filter with 0.22 μm pore size, and loaded onto a 5-mL HisTrap column (Cytiva life sciences) that had been equilibrated in Buffer A (50 mM Tris pH 7.5, 250 mM NaCl, 10 mM imidazole, and 200 μM TCEP). The His-tagged ASCC2 CUE domain that was retained by the HisTrap column was eluted by running a gradient from 0% to 100% Buffer B (50 mM Tris pH 7.5, 250 mM NaCl, 400 mM imidazole, and 200 μM TCEP) over 100 mL. Fractions judged to contain His-tagged ASCC2 CUE domain by gel electrophoresis were combined and incubated with His-tagged TEV protease while being dialyzed in Buffer A overnight at 4°C. The dialyzed sample was then repassed over a HisTrap column equilibrated in Buffer A, and the flowthrough containing the untagged ASCC2 CUE domain was collected and concentrated to less than 5 mL. The concentrated ASCC2 CUE domain solution was passed over a Superdex 75 16/60 size-exclusion column (Cytiva life sciences) equilibrated in 20 mM HEPES pH 7.6, 150 mM NaCl, and 200 μM TCEP. The ASCC2 CUE domain eluted from the column as a single peak roughly 85 mL after injection.

### Expression and purification of full-length ASCC2

*E. coli* Rosetta (DE3) cells were transformed with pET-28 vector containing full-length ASCC2 and grown on LB agar plates with kanamycin and chloramphenicol. The resulting colonies were used to inoculate 5 mL cultures of LB media with kanamycin and chloramphenicol, which were grown overnight at 37°C and 250 rpm shaking until reaching saturation. The 5-mL cultures were used to inoculate 1-L cultures of LB media with kanamycin and chloramphenicol that were grown at 37°C and 250 rpm shaking until reaching an OD_600_ between 0.5 and 0.8. Once the cells had reached the appropriate density, the temperature was lowered to 16°C and ASCC2 expression was induced by adding 500 μL of 1 M IPTG. After approximately 16 hours, the cells were harvested by centrifuging at 5,000 rpm for 20 minutes. The cells were resuspended in 100 mL of lysis buffer (50 mM Tris pH 7.5, 250 mM NaCl, 20 mM imidazole, 3 mM β-mercaptoethanol, 2 μM PMSF, and one cOmplete Mini, EDTA-free protease-inhibitor tablet (Roche)), lysed using a microfluidizer, and then centrifuged at 14,000 rpm for 30 minutes at 4°C to separate the soluble and insoluble fractions. The soluble fraction was then passed through syringe filters with 0.45-micron and 0.22-micron pore sizes prior to being loaded onto a 5-mL HisTrap column (Cytiva life sciences) that had been equilibrated in Buffer A (50 mM Tris pH 7.5, 250 mM NaCl, 20 mM imidazole, 3 mM β-mercaptoethanol). His-tagged, full-length ASCC was eluted from the column using a gradient from 0% to 100% Buffer B (50 mM Tris pH 7.5, 250 mM NaCl, 400 mM imidazole, 3 mM β-mercaptoethanol) over 50 mL. Fractions containing full-length ASCC2, as determined by SDS-PAGE, were concentrated to less than 5 mL total volume and passed over a Superdex 200 16/60 size-exclusion column (Cytiva Life Sciences) that had been equilibrated in a buffer consisting of 20 mM HEPES pH 7.5, 150 mM NaCl, and 200 μM HEPES. Full-length ASCC2 eluted from the Superdex 200 16/60 size-exclusion column 60-70 mL after injection.

### Expression and purification of monoubiquitin

BL21 DE3 *E. coli* cells were transformed with pET-3a vector containing the ubiquitin constructs, plated on LB agar containing 100 μg/mL ampicillin, and incubated overnight at 37°C. Single colonies were used to inoculate 5-mL aliquots of LB media with 100 μg/mL ampicillin. The 5-mL cultures were grown overnight at 37°C with 250 rpm shaking until saturation. The 5-mL colonies were used to inoculate 1-L cultures of LB media with 100 μg/mL ampicillin, which were grown at 37°C with 250 rpm shaking until reaching an OD_600_ between 0.5 and 0.8. Protein expression was induced by adding 0.5 mM IPTG, and allowed to continue overnight at 16°C with 250 rpm shaking. Following protein expression, cell were pelleted by centrifugation, resuspended in a buffer consisting of 50 mM Tris pH 7.5 and 1 mM PMSF, and lysed by sonication on ice. Cell lysate was centrifuged at 17,500 x *g* for 30 minutes at 4°C, after which the soluble fraction was separated and slowly stirred on ice. To the stirring soluble fraction, 1% (v/v) of 70% perchloric acid was added dropwise until the solution turned a milky white. This solution was centrifuged at 17,500 x *g* for 30 minutes at 4°C after which the soluble fraction containing the ubiquitin was separated from the pellet. The soluble fraction was then subjected to multiple rounds of dialysis in 10 mM Tris pH 7.6 until reaching a neutral pH.

### Conjugation and purification of polyubiquitin chains

Polyubiquitin chains were assembled enzymatically by combining monoubiquitin (>1 mM), human UBE1 enzyme (500 nM), and *S. cerevisiae* Ubc13/Mms2 (2.5 μM) in a solution containing 50 mM HEPES pH 7.5, 10 mM MgCl_2_, 1 mM TCEP, and 10 mM ATP. To limit the chain length to diubiquitin, K48R/K63R ubiquitin and D77 ubiquitin can be substituted for wild-type ubiquitin in the reaction mixture (18). Human UBE1 and *S. cerevisiae* Ubc13/Mms2 enzymes were expressed and purified as previously described (19,20). The reaction mixture was incubated overnight at 37°C and then diluted 10-fold in Buffer A (50 mM ammonium acetate pH 4.5 and 50 mM NaCl) and loaded onto a monoS 10/100 GL column (Cytiva life sciences) equilibrated in buffer A. The ubiquitin species retained by the column were eluted by running a gradient from 0-100% buffer B (50 mM ammonium acetate pH 4.5 and 600 mM NaCl) over 300 mL.

### Isothermal titration calorimetry binding experiments and data analysis

For ITC experiments involving K63Ub_2_, distal ubiquitins contained K48R/K63R mutations and proximal ubiquitins contained D77 mutations to control the polyubiquitin chain length, as described above. For the monoubiquitin binding experiment in Figure 2A, K48R/K63R ubiquitin was used. Prior to each isothermal titration calorimetry experiment, proteins were dialyzed overnight in a solution of 20 mM HEPES pH 7.5, 150 mM NaCl, and 200 μM TCEP. Using a MicroCal iTC_200_ instrument (Malvern), titrations were conducted using a series of 2-μL injections each lasting four seconds, with a minimum of two minutes between injections. Fitting was performed using Origin 7 SR4 (OriginLab). To extract *K*_d_ information from non-sigmoidal isotherms, N values were fixed by altering the active concentrations in the cell and syringe during fitting. The first *K*_d_ value in the range corresponds to varying the active concentration in the cell and the second *K*_d_ value in the range corresponds to varying the active concentration in the syringe.

### Chemical shift mapping experiments

^15^N-labeled ASCC2(465-521) and K63-linked diubiquitin were made following similar expression and purification protocols to those described above, however, after reaching an OD_600_ between 0.5 and 0.8, the 1-L aliquots of cells were pelleted, washed with M9 salts, and resuspended in one-third the original volume of minimal media containing ^15^N-labeled ammonium chloride. The resuspended cells recovered for 1 hour at 37°C with 225 rpm shaking prior to proceeding with the induction of protein expression as described above.

^1^H,^15^N-HSQC spectra of ASCC2(465-521) and ubiquitin were recorded using a 600 MHz AVANCE II NMR system at the Biomolecular NMR Center at Johns Hopkins University. Resonances in the ^1^H,^15^N-HSQC spectra of ubiquitin were assigned based on data from Dr. Carlos Castañeda (personal communication). Resonance assignments for the ASCC2(465-521) ^1^H,^15^N-HSQC spectra were obtained from 3D ^15^N-edited ^1^H-^1^H NOESY-HSQC and ^15^N-edited ^1^H-^1^H TOCSY-HSQC spectra. NMR data were processed using nmrPipe software (21) and CSP values were measured using the formula 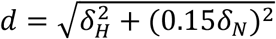 by the program CcpNmr Analysis (22) on the NMRBox platform (23). The CSP values in Figures 3 were measured for the titration of 100 μM K63Ub_2_,^15^N-labeled on the distal ubiquitin in Figure 3A and ^15^N-labeled on the proximal ubiquitin in Figure 3B, with 30 μM, 100 μM, 200 μM, and 500 μM ASCC2(465-521) at 20°C. The CSP values in Figure 5A were measured for 20 μM ^15^N-labeled ASCC2(465-521) alone and in the presence of 681 μM K48R/K63R ubiquitin at 40°C in a buffer consisting of 20 mM Tris pH 7.0, 100 mM NaCl, and 200 μM TCEP. The CSP values in Figure 5B were measured for the titration of 20 μM ^15^N-labeled ASCC2(465-521) with 6 μM, 20 μM, and 40 μM K63Ub_2_ at 40°C in a buffer consisting of 20 mM Tris pH 7.0, 100 mM NaCl, and 200 μM TCEP. In Figures 3 and 5, resonances that disappeared during the course of the titration are marked by an asterisk at the pinnacle of their corresponding bar and assigned values of 0.3 or 0.25 ppm, respectively. NMR data, chemical shift assignments, and CSP values for ^15^N-labeled ASCC2 CUE domain titrated with monoubiquitin and K63Ub_2_ have been deposited to the Biological Magnetic Resonance Data Bank (24) as entries 51130 and 51139, respectively. NMR data, chemical shift assignments, and CSP values for ^15^N-labeled K63Ub_2_ titrated with the ASCC2 CUE domain have been deposited to the Biological Magnetic Resonance Data Bank (24) as entries 51145 (^5^N-labeled on the proximal ubiquitin) and 51146 (^15^N-labeled on the distal ubiquitin).

### Determining ASCC2 CUE domain binding affinity for monoubiquitin using CSP data

The program CcpNmr Analysis (22) used information from the titration of ^15^N-labeled ASCC2(465-521) with monoubiquitin, as described in the previous section, to determine the *K*_d_ value for ASCC2(465-521) binding monoubiquitin using the formula 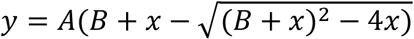 where *y* = d_obs_, *A* = δ_∞_/2, *B* = 1 + *K*_d_/*a, a* = [ASCC2]_tot_, *b* = [Ub]_tot_, *x* = *b*/*a*. The *K*_d_ value of 39.6 μM ± 1.6 μM reported in Supplementary Figure 1 is the average of the *K*_d_ values determined for residues L478, L479, Q500, and L506. These four residues have the largest CSP values recorded for the titration of ^15^N-labeled ASCC2(465-521) with monoubiquitin and are all predicted to be at the ASCC2:ubiquitin binding interface.

### Modeling the ASCC2 CUE:K63Ub_2_ complex using PyMOL and the HADDOCK server

The PyMOL molecular visualization system (25) and the HADDOCK protein-docking server (16,17) were used to model the interaction between the ASCC2 CUE domain and K63Ub_2_. First, the interaction between the ASCC2 CUE domain and the distal ubiquitin was modeled based on the solution structure of the gp78 CUE:monoubiquitin complex (PDB ID: 2LVO) (10). PyMOL was used to superimpose the solution structure of the ASCC2 CUE domain (PDB ID: 2DI0) with the structure of the gp78 CUE domain in the ubiquitin-bound complex. The HADDOCK server was then used to model the interaction between the proximal ubiquitin and the ASCC2 CUE domain. To guide the docking experiment, proximal ubiquitin residues with CSP values greater than 2σ, or resonances that disappeared during the NMR titration, were identified as “active” residues. For the ASCC2 CUE domain, residues outside of the conserved hydrophobic patch that decrease the ubiquitin-binding affinity when mutated were identified as active. For the reported model, the active proximal ubiquitin residues were A46, G47, K48, Q49, E64, T66, and L71, and the active ASCC2 CUE domain residues were E467 and S470. Additionally, a distance restraint of 1.32 Å between the carbonyl carbon of G76 of the distal ubiquitin and the ε-amino group of K63 of the proximal ubiquitin was used to approximate the isopeptide linkage in the diubiquitin chain. The reported model is the best structure from the highest scoring cluster.

### Immunofluorescence analysis of HA-tagged ASCC2

All immunofluorescence analysis was performed in U2OS cells using wild-type and mutant forms of ASCC2, expressed in the pHAGE-CMV-3xHA lentivirus (7). Three days after transduction, the cells were treated with 500 μM MMS in complete DMEM media for six hours. U2OS cells were washed once with ice-cold PBS, then extracted with 1× PBS containing 0.2% Triton X-100 and protease inhibitors (Pierce) for 10–20 min on ice before fixation with 3.2% paraformaldehyde. The cells were then washed extensively with immunofluorescence wash buffer (1× PBS, 0.5% NP-40, and 0.02% NaN_3_), then blocked with immunofluorescence blocking buffer (immuno-fluorescence wash buffer plus 10% FBS) for at least 30 min. Primary antibodies were diluted in immunofluorescence blocking buffer overnight at 4 °C. After staining with secondary antibodies (conjugated with Alexa Fluor 488 or 594; Millipore) and Hoechst 33342 (Sigma-Aldrich), where indicated, samples were mounted using Prolong Gold mounting medium (Invitrogen). Epifluorescence microscopy was performed on an Olympus fluorescence microscope (BX-53) using an ApoN 60×/1.49 numerical aperture oil immersion lens or an UPlanS-Apo 100×/1.4 numerical aperture oil immersion lens and cellSens Dimension software. Raw images were exported into Adobe Photoshop, and for any adjustments in image contrast or brightness, the levels function was applied. For foci quantification, at least 100 cells were analyzed in triplicate.

## Data Availability

NMR data reported in this manuscript have been deposited in the Biological Magnetic Resonance Data Bank as entries 51130 (^15^N-labeled ASCC2 CUE domain interacting with monoubiquitin), 51139 (^15^N-labeled ASCC2 CUE domain interacting with K63Ub_2_), 51145 (K63Ub_2_ ^15^N-labeled on the proximal ubiquitin interacting with the ASCC2 CUE domain), and 51146 (K63Ub_2_ ^15^N-labeled on the distal ubiquitin interacting with the ASCC2 CUE domain).

## Acknowledgements

The authors acknowledge Dr. Stoyan Milev for his assistance interpreting ITC data with low C values and Drs. Aswani Kumar Kancherla and Dominique Frueh for their help processing and analyzing NMR data. The authors also thank Dr. Carlos Castañeda for providing the assignments from a previously recorded ^1^H,^15^N-HSQC ubiquitin spectrum that served as a guide for the ubiquitin assignments in this manuscript.

## Funding and additional information

Research reported in this publication was supported by National Institute of General Medical Sciences awards GM140410 (P.M.L.) and GM130393 (C.W.), National Cancer Institute awards CA227001 and CA092584 (N.M.), and American Cancer Society research scholar grant RSG-18-156-01-DMC (N.M.). J.G.B and L.N.G were supported by NSF S-STEM Award 1458490. This study made use of NMRbox: National Center for Biomolecular NMR Data Processing and Analysis, a Biomedical Technology Research Resource (BTRR), which is supported by the National Institute of General Medical Sciences (GM111135). The content is solely the responsibility of the authors and does not necessarily represent the official views of the National Institutes of Health.

## Abbreviations and nomenclature

ALKBH3: Alpha-ketoglutarate-dependent dioxygenase alkB homolog 3
ASCC2: Activating Signal Cointegrator 1 Complex Subunit 2
ASCC3: Activating Signal Cointegrator 1 Complex Subunit 3
ATP: adenosine triphosphate
BBR2: pre-mRNA splicing helicase BRR2
CSP: chemical shift perturbation
CUE: coupling of ubiquitin conjugation to ER degradation
DTT: dithiothreitol
HADDOCK: High Ambiguity Driven protein–protein DOCKing
HSQC: Heteronuclear Single Quantum Coherence
ITC: isothermal titration calorimetry
PMSF: phenylmethylsulfonyl fluoride
ppm: parts per million
RNF113A: ring finger protein RNF113A
RMSD: root-mean-square deviation
rpm: revolutions per minute
TCEP: Tris (2-carboxyethyl) phosphine
TEV: tobacco etch virus

