## Supplemental Figures for "The ASCC2 CUE domain contacts adjacent ubiquitins to recognize K63-linked polyubiquitin"

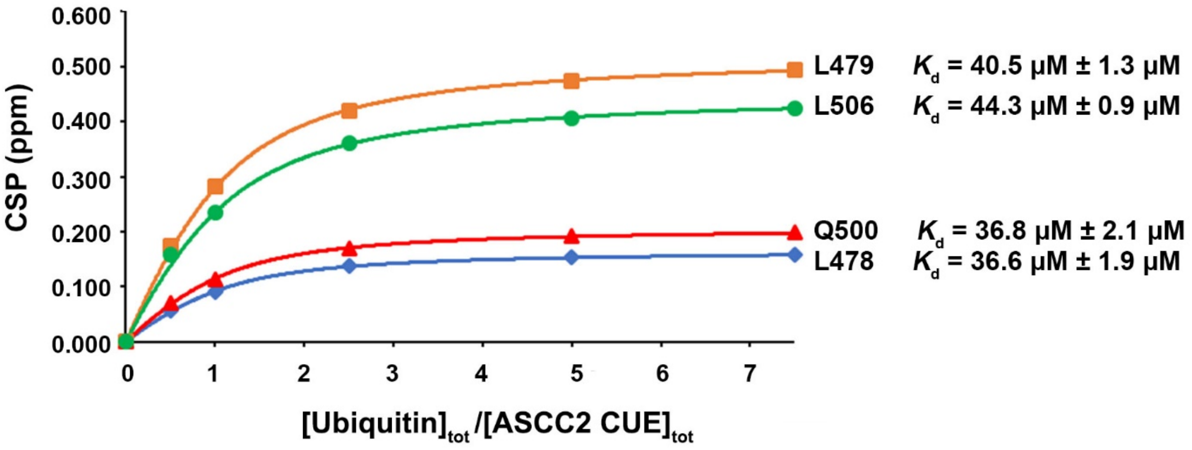


Supplementary Figure 1. Determination of ASCC2 CUE domain binding affinity for monoubiquitin using NMR spectroscopy. $^{1}H,^{15}N$-HSQC spectra of 100 μM $^{15}N$-labeled ASCC2 CUE domain were recorded alone and in the presence of 50, 100, 250, 500, and 750 μM monoubiquitin. The chemical shift perturbations (CSPs) for ASCC2 CUE domain residues L478, L479, Q500, and L506 at the ubiquitin-binding interface were used to determine the binding affinity for monoubiquitin. The average of the $K_d$ values for residues L478, L479, Q500, and L506 is 39.6 μM ± 1.6 μM.

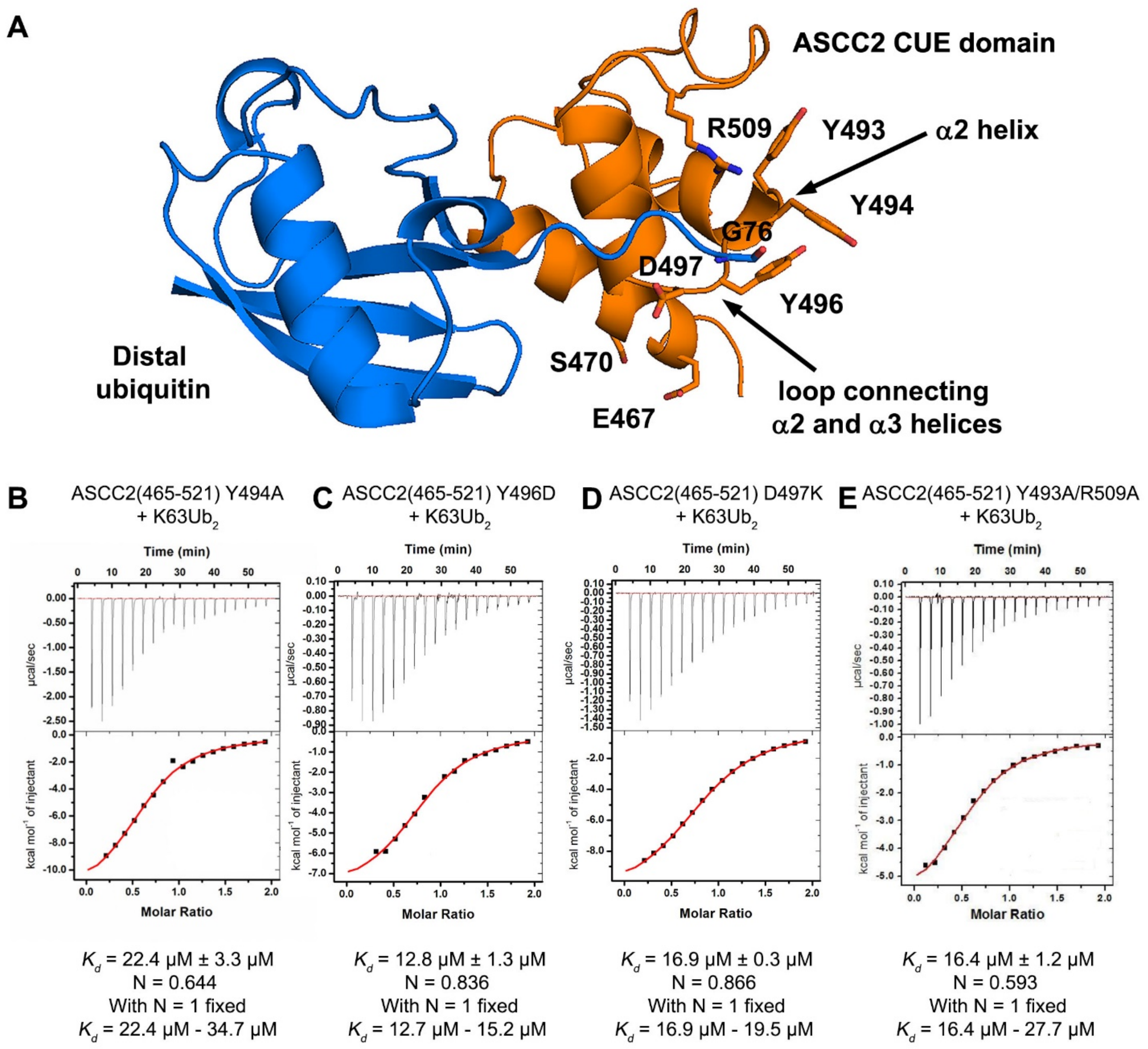


Supplementary Figure 2. Probing for ASCC2 CUE domain residues that contact the proximal ubiquitin of K63Ub$_2$. (A) Model of the distal ubiquitin of K63Ub$_2$ bound to the ASCC2 CUE domain based on the structure of the gp78 CUE:ubiquitin complex (PDB ID: 2LVO). Given the positioning of the C-terminal tail of the distal ubiquitin, residues from the α2 helix, or the loop connecting the α2 and α3 helices, could be close enough to contact the proximal ubiquitin. However, Y494A (B), Y496D (C), D497K (D), and Y493A/R509A (E) ASCC2 CUE domain mutants bind K63Ub$_2$ with similar affinity to the wild-type ASCC2 CUE domain, suggesting these residues do not contact the proximal ubiquitin of K63Ub$_2$.

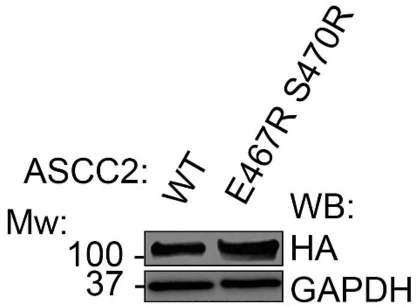


Supplementary Figure 3. Comparison of ASCC2 expression levels in Figure 6E. U2OS cells were transduced with WT HA-ASCC2 or the E467R/S470R mutant. Whole cell lysates were collected and used for western blotting using anti-HA antibody. GAPDH was used as the loading control.
